# Neural Mechanisms of tDCS: Insights from an In-Vivo Rodent Model with Realistic Electric Field Strengths

**DOI:** 10.1101/2024.12.20.629687

**Authors:** Liyi Chen, Alireza Majdi, Boateng Asamoah, Myles Mc Laughlin

## Abstract

**Introduction:** Transcranial direct current stimulation (tDCS) is a non-invasive neuromodulation method using low amplitude current (1-2 mA) to create weak electric fields (<1 V/m) in the brain, influencing cognition, motor skills, and behavior. However, the neural mechanisms remain unclear, as prior studies used high electric field strengths (10–40 V/m) unrepresentative of human tDCS.

**Objective:** This study aimed to develop an in-vivo rat model replicating human tDCS electric field strengths to examine effects of weak electric fields on cortical neurons.

**Method:** Currents of 0.005–0.3 mA were applied in 9 rats, generating electric fields of 0.5–35 V/m in the somatosensory cortex. Neural activity across cortical layers was recorded using a multichannel silicone probe. Somatosensory evoked potentials (SSEP) elicited by foot shocks assessed membrane polarization. Regular spiking (RS) and fast-spiking (FS) neurons were identified via spike shapes. Effects of tDCS on SSEP, spontaneous spiking activity (SSA), and evoked spiking activity (ESA) were analyzed.

**Results:** Anodal tDCS caused hyperpolarization (SSEP increase) in superficial layers and depolarization (SSEP decrease) in deeper layers, reversing asymmetrically for cathodal stimulation. Weak fields (<1 V/m) altered SSA in RS but not FS neurons, while stronger fields affected ESA in RS neurons. Effects correlated with field strength and were well described by linear mixed-effect models. Changes in SSA were correlated with changes in SSEP.

**Conclusion:** This study demonstrates that realistic tDCS fields induce complex cortical polarization patterns linked to SSA changes. Increasing electric field strength amplifies effects, suggesting higher amplitude tDCS could enhance efficacy in humans.

**Highlights:**

1. Newly developed in-vivo rodent model to replicate the weak electric field strengths characteristic of human tDCS, probe localized membrane polarization effects and simultaneously monitor spontaneous and evoked spiking activity.
2. Results provide the first direct in-vivo confirmation of several tDCS mechanistic predictions derived from computational models and brain slice work.
3. Complex Membrane Polarization Patterns: tDCS induces simultaneous hyperpolarization and depolarization in distinct neuronal compartments.
4. Neuron-Specific Effects: Weak electric fields preferentially modulate excitatory neurons, with no significant impact on inhibitory neurons at low electric field strengths.
5. Polarity asymmetric effects: Anodic stimulation produces stronger effects than cathodic stimulation.
6. Membrane polarization is linked to changes in spiking activity: Changes in membrane polarization are correlated with changes in spontaneous spiking activity.
7. All the tDCS neural mechanisms showed effects that were linearly related to electric field strength, underscoring the translational importance of novel tDCS protocols that can increase electric field strength, potentially improving the robustness and reproducibility of tDCS protocols in humans.

## 1. Introduction

Transcranial direct current stimulation (tDCS) is a non-invasive neuromodulation method in which low amplitude current (1-2 mA) is passed through scalp electrodes to create a weak electric field (<1V/m) in the brain ^1^. Numerous studies in both healthy volunteers and patients have shown how tDCS can cause a wide range of effects on cognition, motor skill and behavior ^2–4^. However, recent robust studies have failed to reproduce some of these key results ^5–8^. One key issue in the field is that the effects of weak electric fields on cortical neurons (i.e. the proposed mechanism through which tDCS works) remain poorly understood ^9^.

Most work on understanding the neural mechanisms underpinning tDCS effects in humans has focused on applying uniform DC stimulation to brain slices ^10–13^.These important studies have proposed several distinct and often complex neural mechanisms through which tDCS may work. In practice, these distinct tDCS mechanisms may interact with one another to cause the tDCS effects observed in humans. Below we highlight the most important tDCS mechanisms, together with the current limitations in their understanding.

The rationale behind many human tDCS studies typically follows the ‘somatic doctrine’ ^14–16^ where anodal tDCS is assumed to cause cortical excitation, while cathodal tDCS causes cortical inhibition. However, work in slices indicates a much more complex pattern of effects where DC stimulation causes hyperpolarization and depolarization to occur simultaneously across different neuronal compartments ^17–19^. Slice work has shown that neuronal morphology also strongly affects the degree of membrane polarization with excitatory pyramidal neurons exhibiting much stronger polarization effects than inhibitory interneurons ^18,20,21^. Another level of complexity is that while anodic and cathodic DC stimulation have been shown to have opposite effects on membrane polarization, there is an asymmetry in potency, with anodic stimulation showing stronger effects in slices ^19^. The effects of tDCS on membrane polarization are typically not strong enough to directly trigger action potentials ^22^. However, tDCS has been shown to cause changes in spontaneous spiking activity (SSA) in animal models^23–26^. While the existing works have advanced our understanding of tDCS neural mechanisms, two key outstanding limitations remain: First, most tDCS animal and slice work has used relatively high electric field strengths (10 to 40 V/m), which are never achieved in humans tDCS. This implies that the potential role of the aforementioned mechanisms in producing the tDCS effects observed in humans remains a topic of debate ^27^. Second, a link between two of these tDCS mechanisms has been proposed, but never experimentally demonstrated: namely that the complex changes in membrane polarization led to a change in SSA.

To help answer these gaps in our understanding, we created an in-vivo tDCS rat model with electric field strengths matching human tDCS and used it to study tDCS mechanisms. We used somatosensory evoked potentials (SSEPs) sampled across cortical layers as a proxy measure for membrane polarization of different neural compartments. Our results confirmed that anodic stimulation caused hyperpolarization of the apical dendrites and depolarization of the soma and basal dendrites. Cathodic stimulation caused an asymmetric reversal of this pattern. Next, we made recordings of spiking activity and separated these into putative excitatory and inhibitory neurons. We found weak electric fields of <1V/m had a significant effect on the SSA of putative excitatory neurons, but not on putative inhibitory neurons. Strong electric fields were needed to influence evoked spiking activity. Finally, changes in membrane polarization across different neural compartments were found to be linearly correlated with changes in SSA. Our results help resolve two important limitations in our current understanding of how tDCS works: First, we showed that many of the tDCS mechanisms demonstrated in brain slices are present in-vivo at electric field strengths achieved in humans tDCS. Importantly these effects become stronger with increasing electric field strength. Second, we present the first experimental evidence linking the complex pattern of membrane polarization to changes in SSA. This work advances our current understanding of tDCS mechanisms, and its application in human studies has the potential to enhance the robustness of tDCS effects.

## 2. Result

### 2.1 Quantification of the relationship between stimulation current and cortical electric-field strength

The strength of the electric field (E-field) generated in the cortex by tDCS is a key factor affecting both evoked potentials and neural spiking activity. Therefore, it is essential to measure and calibrate the cortical E-field generated when different tDCS amplitudes are applied. To do this, we used a 32-channel silicon probe (50 µm pitch, spanning 1550 µm) to record voltage (i.e. stimulation artifact) across different layers in the somatosensory cortex using a 100Hz sinewave at four different stimulation amplitudes (0.01, 0.015, 0.02, and 0.025 mA). Similar to previous work ^28^, voltage decreased across cortical depths with some fluctuations and scaled linearly with stimulation amplitude (Fig. 1A, example recording from one rat). The voltage data collected in eight rats were pooled and fit with a linear mixed effects model, with the slope of the linear model fit across the different voltages. Fig. 1B shows the model-fitted voltage data with solid lines representing the model-estimated voltage and the shaded areas indicating the 95% confidence intervals. ANOVA of the linear-mixed model showed a significant effect of stimulation amplitude on voltage (Fstat=120.7, p<0.001) and a significant interaction between stimulation amplitude and cortical depth (Fstat=10.563, p<0.001) but cortical depth effect was not significant (Fstat=0.001, p=1). We then calculated the E-field by taking the spatial derivative of the modeled voltage data, providing a detailed map of the E-field distribution across cortical depth (Fig. 1C). To illustrate the relationship between tDCS amplitude and E-field strength, we plotted the model-fitted E-field data for five example depths (200μm, 600μm, 850μm, 1250μm, and 1500μm) and then linearly extended the modeled data to show the relevant tDCS amplitudes and E-fields used in all other experiments (Fig.1D). These data were used in the following results to relate the applied tDCS amplitude to an E-field at each cortical depth.

**Figure 1.**
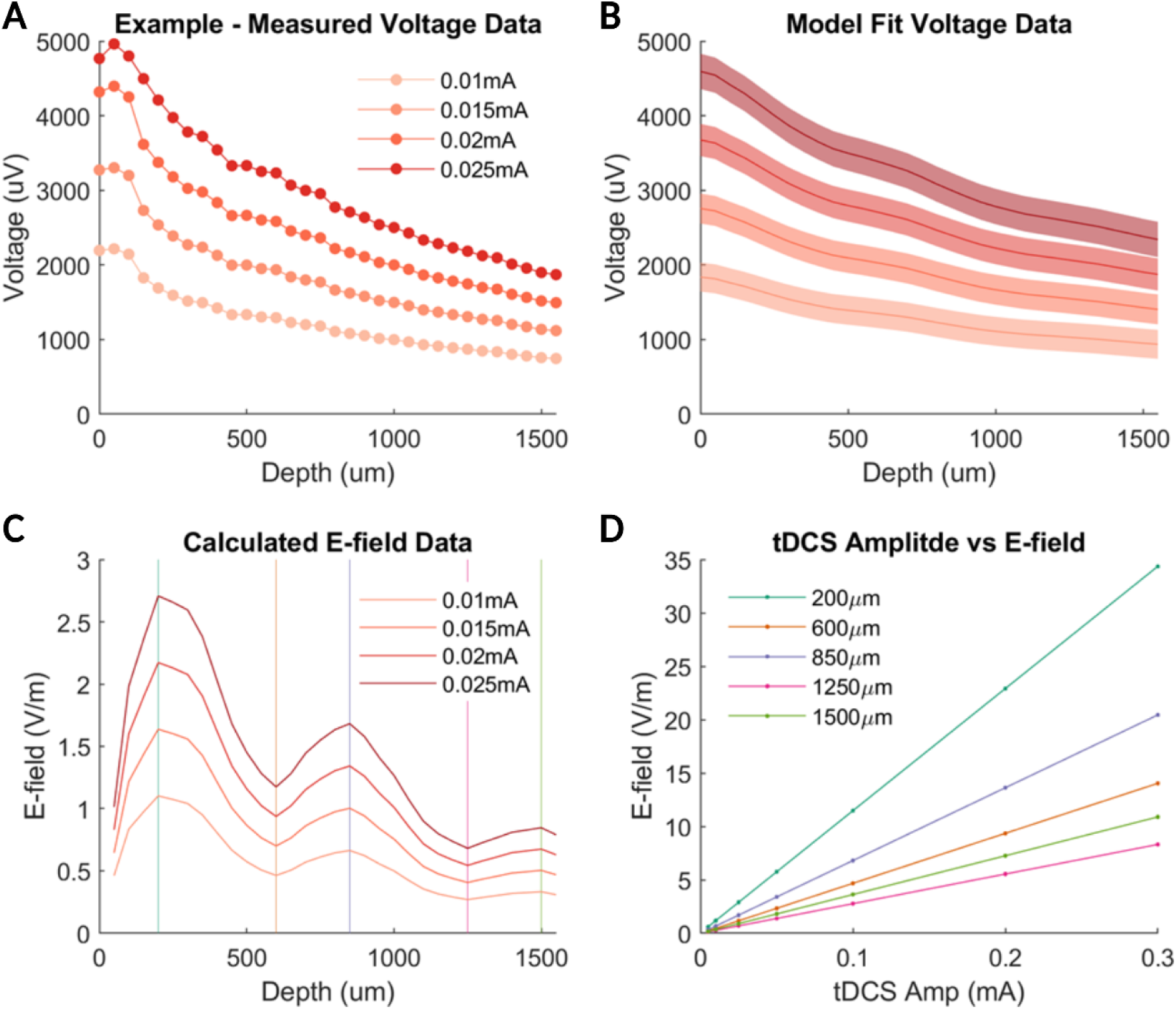
Measurement and statistical modeling of the electric field (E-field). A) The voltage data were measured along one electrode track in the somatosensory cortex when 4 different stimulation amplitudes were applied (0.01, 0.015, 0.02, and 0.025 mA). B) Voltage data from all rats (n=8) were fit with a linear mixed-effects model. The means (solid lines) and 95% confidence intervals of the modeled data are shown. C) The E-field component in the direction of the electrode (i.e. cortical depth) was then calculated by taking the spatial derivative of the modeled data. D) The E-field data shown in C for 5 example depths are replotted as a function of tDCS amplitude vs E-field strength. The modeled data are extended to show the relevant tDCS amplitudes and E-fields used in all other experiments in the paper. Colored lines in panel D correspond to the vertical-colored lines in panel C.

### 2.2 tDCS caused a polarity-dependent pattern of simultaneous increases and decreases in SSEP amplitude across different cortical layers

We used a 32-channel silicon probe to measure SSEPs (elicited via forepaw shock) across different layers in the somatosensory cortex. Fig. 2A shows a cartoon of a typical layer 4 and 5 pyramidal neuron and how it is positioned across different cortical layers, together with the E-field data from Fig. 1. Fig. 2B shows the baseline SSEP (grey lines) and the SSEP recorded during +0.3 mA tDCS (red lines). Compared to baseline, anodic tDCS resulted in larger SSEPs in superficial cortical layers and smaller SSEPs in deeper cortical layers. To quantify SSEP changes, we calculated the difference between baseline and tDCS SSEPs (ΔSSEP = SSEP_tDCS_ – SSEP_baseline_), encoded as color with time on the x-axis (Fig. 2C). This analysis revealed a complex spatiotemporal pattern, showing a localized decrease in ΔSSEP (hyperpolarization) near the cortical surface and localized increase in ΔSSEP (depolarization) in deeper regions. We further quantified this by plotting the ΔSSEP at the SSEP trough (∼10ms) as a function of cortical depth, highlighting the specific depths at which these changes occur (Fig.2D). We did the same analyses for -0.3 mA tDCS (Fig. 2E, 2F, and 2G). The results show that the complex ΔSSEP spatiotemporal pattern is generally reversed under this condition. However, there is an asymmetry in the response to anodal and cathodal tDCS with the switch between localized increase and decrease in ΔSSEP occurring at different depths. The pattern of anodal tDCS causing larger SSEPs in superficial and smaller SSEPs in deeper cortical layers, and the asymmetrical reversal of the pattern upon tDCS polarity reversal matches closely the complex pattern of simultaneous localized hyperpolarization and depolarization of different neural compartments observed in brain slice work when DC fields are applied ^10,29^. Thus, here we propose that ΔSSEP can serve as a proxy metric reflecting localized changes in membrane polarization caused by tDCS.

**Figure 2.**
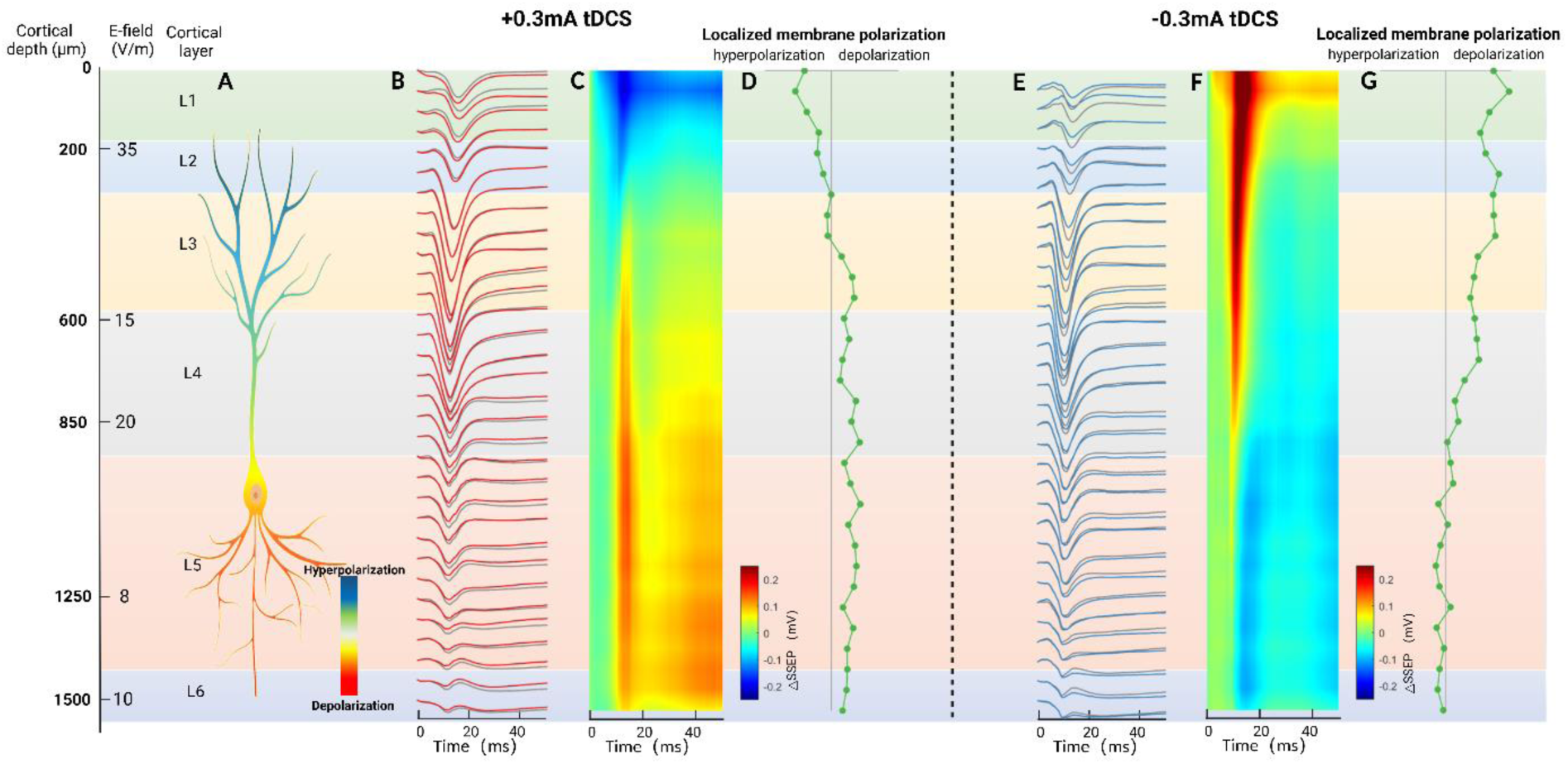
Individual example showing the complex spatiotemporal effects of tDCS on localized membrane polarization quantified via changes in somatosensory evoked potential (SSEP). Far left shows cortical depth, layer, model estimated E-field, and hypothesized effects on a layer 5 pyramidal neuron. A) Example SSEP measured at baseline (SSEPbaseline, grey) and during 0.3mA tDCS (SSEPtDCS, red) at different cortical depths. The X-axis shows time. B) Shows the difference in the baseline and tDCS SSEPs (△SSEP = SSEPtDCS – SSEPbaseline) encoded as color with time on the x-axis. This analysis reveals a complex spatiotemporal pattern with anodic tDCS causing a localized decrease in excitability close to the cortical surface accompanied by a localized increase in cortical excitability at deeper cortical regions. C) The maximum localized membrane polarization change (occurring around 10 ms) is plotted as a function of cortical depth. D), E), and F) Show the same but for -0.3 mA tDCS. The complex spatiotemporal pattern of localized membrane polarization is reversed with cathodic tDCS.

For each of the nine rats, we then found the minimum of the negative trough (∼10ms) and calculated △SSEP_n10_ (SSEP_tDCS_n10_ – SSEP_baseline_n10_) at that time point. Fig. 3A shows this data for one rat across different tDCS amplitudes. The △SSEP_n10_ data from all nine rats were then grouped and fit with a linear mixed effects model, with the linear fit applied across the different tDCS amplitudes (i.e. tDCS amplitude was used to predict △SSEP_n10_). Fig. 3B shows the model-fit data with solid lines representing the model estimated △SSEP_n10_ and the shaded areas indicating the 95% confidence intervals. The same data is replotted in Fig. 3C to highlight the linear effect of tDCS amplitude (and thus E-field) on △SSEP _n10_ across different cortical depths. ANOVA of the model fit showed a significant effect of anodic tDCS amplitude on △ SSEP_n10_ (Fstat=784.87, p<0.001), a significant effect of cortical depth (Fstat=4.1003, p=0.043), and a significant interaction between anodic tDCS amplitude and cortical depth (Fstat=535.7, p<0.001). Thus, anodic tDCS led to an increase in localized membrane polarization at the superficial layers - defined as the SSEP becoming less negative compared to baseline, while at deeper layers anodic tDCS had the opposite effect on localized membrane polarization. This pattern increased linearly with increasing anodal tDCS amplitude.

**Figure 3.**
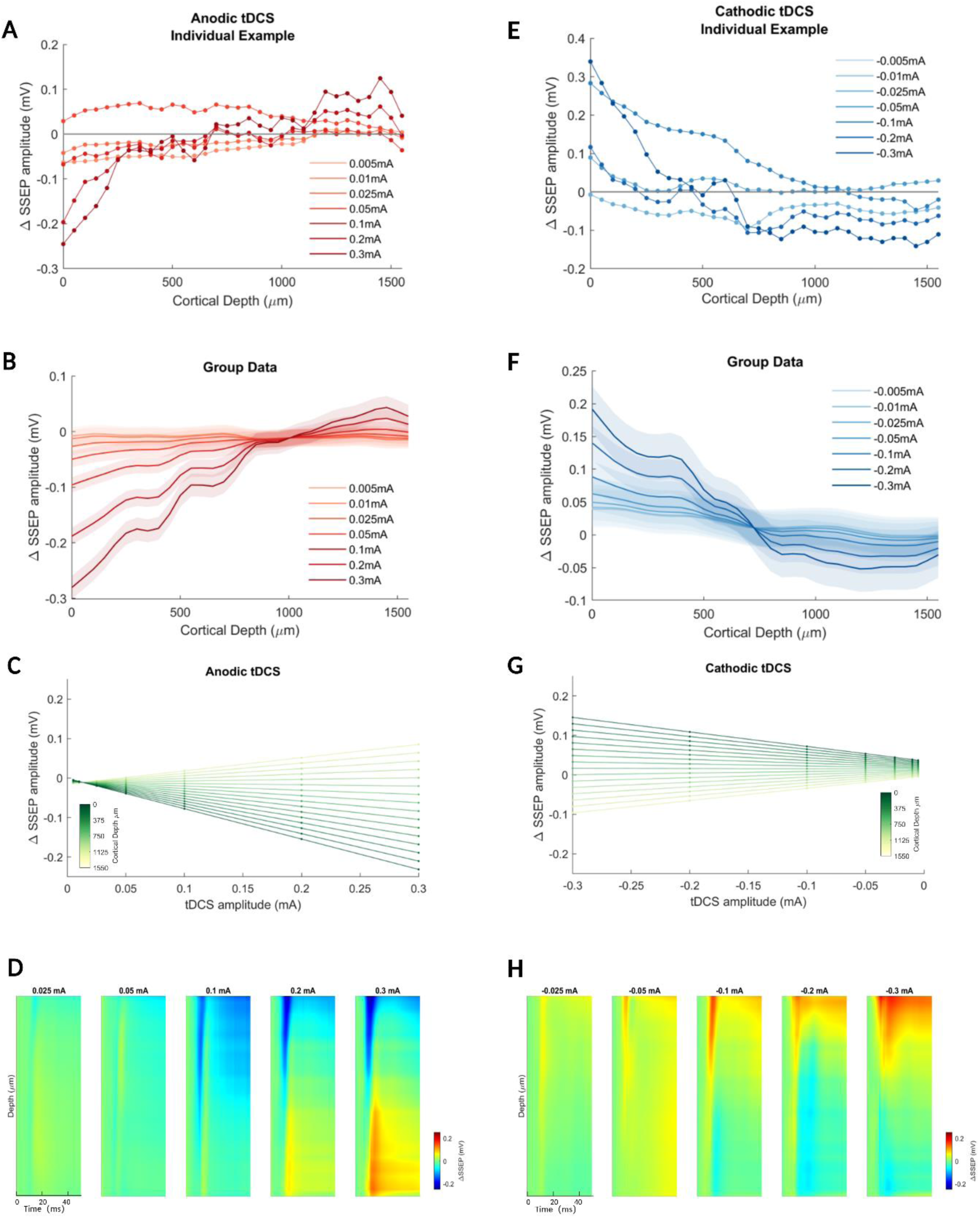
Individual examples and statistical modeling showing the effect of tDCS amplitude and polarity on localized membrane polarization. A) Individual example showing the effect of different amplitudes of anodic tDCS on△SSEP at different cortical depths. B) This data for all rats (n=9) was fit with a linear mixed-effects model. The means (solid lines) and 95% confidence intervals of the modeled-fitted data are shown. C) The model fit from B is replotted to show the linear effect of tDCS amplitude on △SSEP at different cortical depths. D) Anodic tDCS caused an increase in localized membrane polarization (defined as the SSEP becoming more negative during tDCS compared to baseline) at the cortical surface accompanied by a decrease in excitability (defined as the SSEP becoming less negative) at deeper cortical regions. This pattern became stronger as anodal tDCS amplitude was increased. E-H) Show the same for cathodic tDCS which caused the opposite effect.

To gain more insight into the temporal aspects of the effect of tDCS on the SSEP we first calculated the average baseline SSEP for all nine rats (SSEP_baseline_mean_) and did the same for the SSEP during anodic tDCS (SSEP_tDCS_mean_). We then subtracted these to calculate the average change in SSEP (SSEP_mean_ = SSEP_tDCS_mean_ - SSEP_baseline_mean_). These data are shown in Fig 3D for all anodic tDCS amplitudes with time on the x-axis, cortical depth on the y-axis and color representing △SSEP_mean_ . Here we see the effects of anodic tDCS on average change in SSEP peak at around 10ms before slowly decreasing.

The same analysis was applied to the SSEP data in response to cathodic tDCS (Fig 3E-H). ANOVA of the linear-mixed model fit showed a significant effect of cathodic tDCS amplitude △SSEP_n10_ (Fstat=60.499, p<0.001), a significant effect of cortical depth (Fstat=8.5083, p=0.00359), and a significant interaction between cathodic tDCS amplitude and cortical depth (Fstat=66.473, p<0.001). Cathodic tDCS caused the opposite effect compared to anodic tDCS. However, one important difference was that the switch from localized excitability to localized inhibition occurred at a more superficial cortical layer for cathodic tDCS compared to the anodic tDCS. Anodic tDCS also gave a larger absolute △SSEP.

### 2.3 Current source density analysis reveals the effect of tDCS on cortical extra-cellular current flow

Current source density (CSD) analysis can be used to gain insight into the current sinks and sources that generate the SSEP. To perform CSD we first averaged all the baseline SSEPs for nine rats and averaged the SSEPs for each tDCS amplitude condition. Fig. 4A and B show this data for +0.3 mA tDCS. We then calculated the CSD_baseline_ and CSD_tDCS_ by taking the second spatial derivative of the SSEP_baseline_ and SSEP_tDCS_ (Fig. 4C and D) respectively. Both CSD_baseline_ and CSD_tDCS_ show a similar pattern of a sink located in layers 3 and 4 and two sources located in layers 1 and 5. This is consistent with the known anatomy of layer 4 and 5 cortical neurons which receive dense synaptic input in layers 3 and 4 from the ventral posteromedial nucleus (see cartoon in Fig. 4). To quantify the effect of anodic tDCS on extra-cellular current flow we then calculated the △CSD (△CSD= CSD_tDCS_ – CSD_baseline_, see Fig. 4E). This analysis shows that tDCS appears to affect current flow across both superficial and deep cortical layers. Fig. 4F-J show the same analyses for -0.3mA tDCS.

**Figure 4.**
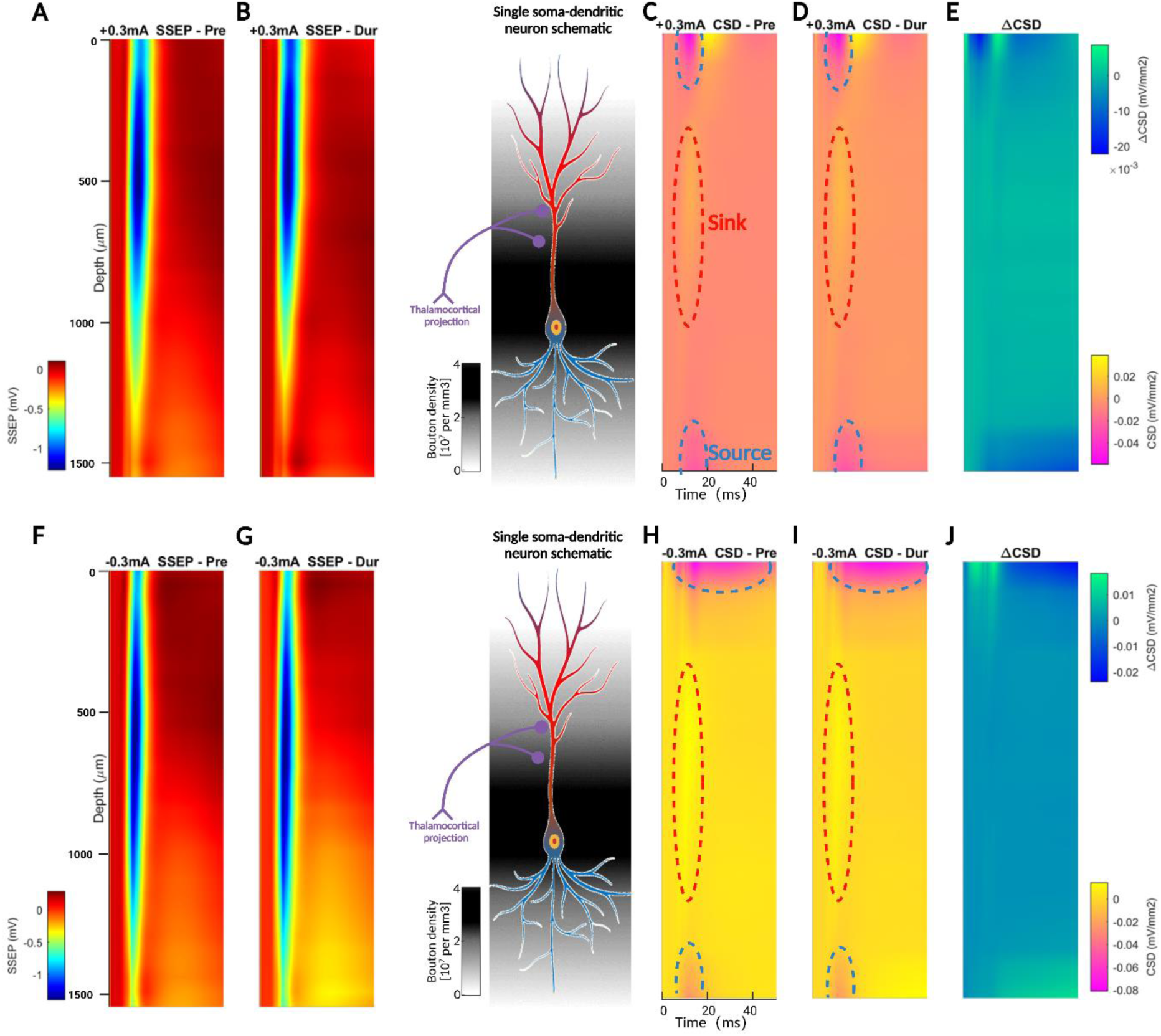
The effect of anodic and cathodic tDCS on the SSEP and related current source density (CSD) analysis. A) Shows the SSEP in response to foot shock as a function of time (x-axis) across different cortical depths (y-axis). B) Shows the same SSEP collected during +0.3 mA tDCS. C) Shows the CSD calculated from the SSEP in panel A. D) Shows the CSD calculated from the SSEP in panel B during +0.3 mA tDCS. Note on both C and D that the current sink in layers 3 and 4 at around 10 ms is accompanied by both a deeper and more superficial source. E) Shows the ΔCSD calculated by subtracting the data in panel C from the data in panel D. This analysis highlights how the application of tDCS can affect extra-cellular current flow in the cortex. F-J) Shows the same for -0.3mA tDCS. The cartoon in the middle shows a typical layer 4 and 5 cortical neurons with button density for ventral posteromedial nucleus inputs^30^.

### 2.4 tDCS effects on spontaneous spike activity

To quantify the effects of tDCS on spiking activity we performed spike sorting on the 32-channel silicon probe data, identifying a total of 99 single neurons. To further assess the effects of tDCS on different neuron types we classified these neurons into regular spiking (RS, n=81) neurons and fast-spiking (FS, n=18) neurons based on the spike waveforms (middle panel of Fig. 5C). As shown in Fig. 5A, RS neurons, with trough-to-peak latency (AP duration) greater than 0.45 ms, are typically assumed to be excitatory neurons, while FS neurons, with AP duration less than 0.45 ms, are considered inhibitory neurons ^31^. Fig. 5A shows a histogram of the AP duration with RS neurons colored yellow and FS green. For each neuron, we then separated spikes into evoked activity occurring within a 50ms time window beginning after the foot shock used to elicit the SSEP and spontaneous activity defined as spikes occurring at all other times. The effect of tDCS on evoked spiking activity is analyzed in section 2.6, the rest of this section focuses on SSA.

**Figure 5.**
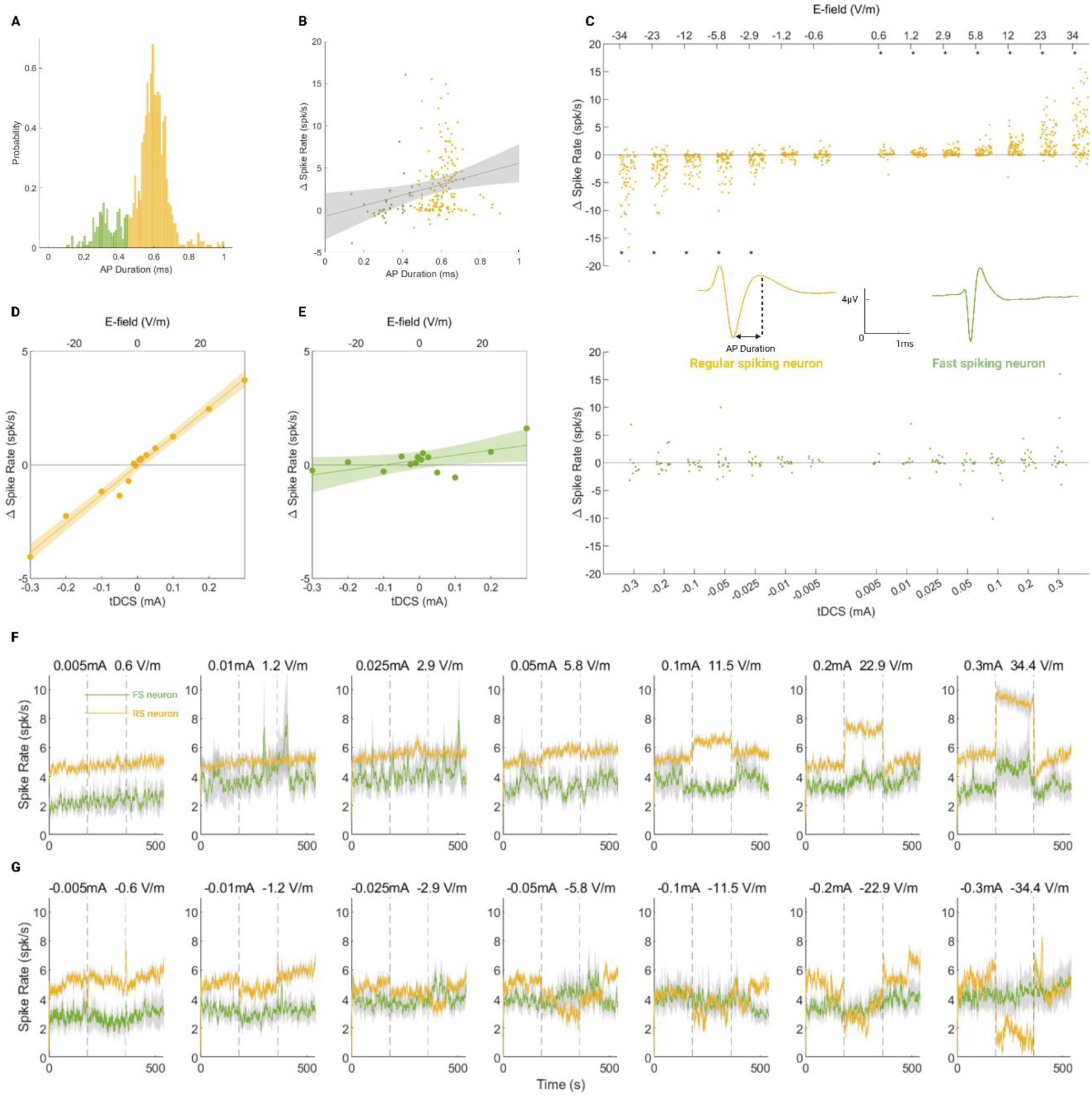
Effect of tDCS amplitude and polarity on spontaneous single unit activity. A) Single neurons were separated into fast-spiking (FS, green) and regular-spiking (RS, yellow) neurons based on action potential duration (see inset in F and G). B) The change in spike rate is plotted as a function of action potential duration for both FS and RS neurons. The dashed line and grey error bars show a linear mixed model fit (n = 97). C) Individual single-neuron data showing how different amplitudes of both anodic and cathodic tDCS caused changes in spike rate (Δ spike rate). For RS neurons cathodic tDCS caused a decrease in spike rate while anodic tDCS caused an increase in spike rate. For FS neurons these effects were mostly absent. The grey dots represent significant changes. D and E) Show the same data as in C but now fit with a linear mixed effects model, with each dot representing the mean Δ spike rate for that condition. F and G) Show how tDCS amplitude and polarity affect the average spike rate over time for both FS and RS neurons.

Fig. 5B plots the change in spontaneous spike rate (ΔSSA) caused by +0.3 mA tDCS as a function of action potential duration for both FS and RS neurons. Data from all neurons were fit with a linear mix- effects model to determine if there was a relationship between AP duration and change in spike rate due to tDCS. The dashed line and grey shadow represent a linear mixed model fit with 95% confidence intervals. The model showed a significant effect of AP duration on Δ spike rate (Fstat=6.9905, p=0.009), i.e. neurons with a longer AP duration show larger changes in spike rate due to tDCS.

Fig. 5C shows ΔSSA as a function of tDCS amplitude for all RS and FS neurons. We found that for RS neurons, cathodic tDCS caused a decrease in SSA, whereas anodic tDCS caused an increase. For FS neurons, these effects were absent. The data was fit with a linear mixed-effects model. We found that tDCS amplitude (Fstat=707.28, p<0.001) had a significant effect on ΔSSA. We also found a significant interaction between tDCS amplitude and neuron type (RS or FS) (Fstat=76.736, p<0.001) but neuron type alone did not have a significant effect on ΔSSA (Fstat=1.8909 p=0.169). Post hoc testing showed that for RS neurons at all anodic tDCS amplitudes ΔSSA were significantly higher than 0: +0.005mA (z=2.98, p=0.040), 0.01mA (z=4.24, p<0.001), 0.025mA (z=5.20, p<0.001), 0.05mA (z=6.26, p< 0.001), 0.1mA (z=6.91, p<0.001), 0.2mA (z=7.32, p<0.001), 0.3mA (z=6.93, p<0.001). Cathodic tDCS conditions at -0.025mA (z=-4.50, p<0.001), -0.05mA (z=-5.63, p<0.001), -0.1mA (z=-6.37, p<0.001), -0.2mA (z=-6.33, p<0.001) and -0.3mA (z=-6.93, p<0.001) were significantly higher than 0, however, ΔSSA were not significantly different in the conditions at -0.005mA (z=-0.42, p=9.438) and -0.01mA (z=2.33, p=0.275). For FS neurons, post-hoc testing showed that none of the anodic nor cathodic tDCS amplitudes had a significant effect on ΔSSA. These results show weak electric fields, comparable to those found in the human cortex during tDCS, can have a significant effect on SSA.

Fig5. D and E show the linear mixed-effects model fit (solid line) and 95% confidence intervals (shaded) for both RS and FS neuron types respectively (model analysis reported above in Fig. 5C). The dots represent the mean ΔSSA data for each tDCS amplitude. Fig.5F shows the average spike rate over time for both RS and FS neurons across all anodic tDCS amplitudes tested. Fig. 5G shows the same data for cathodic tDCS.

### 2.5 tDCS changes in SSEP amplitude are related to changes in spontaneous spike activity

We have shown, via a proxy ΔSSEP metric, that tDCS causes a complex pattern of simultaneous membrane hyperpolarization and depolarization (see Fig. 2 and 3). Additionally, we have shown that tDCS causes changes in SSA (see Fig. 5). Both of these mechanisms were found to: 1) scale linearly with electric field strength (Fig. 2C, G and Fig. 4D), 2) reverse upon tDCS polarity reversal, 3) be stronger for anodic than cathodic tDCS and 4) be present at electric field strengths relevant for human tDCS. Therefore, we next wanted to investigate the relationship between these two tDCS mechanisms.

To do this we first analyzed the effects of tDCS on Δ SSA across different cortical depths and for different neuron types. Fig. 6A shows a histogram distribution for both RS and FS neuron types binned across different cortical depths. The RS neurons sampled were mostly positioned in layers 4 and 5, while the FS neurons sample were spread approximately equally across all layers. Fig. 6B shows the mean changes in SSA due to either +0.3 mA or -0.3 mA tDCS for each neuron type binned across different depths. The largest changes in the spike rate occurred for RS neurons in layers 4 and 5, with neurons in either deeper or more superficial layers showing smaller changes in spike rate. A similar pattern appeared to be visible for FS neurons, but the effect on ΔSSA was lower.

**Figure 6.**
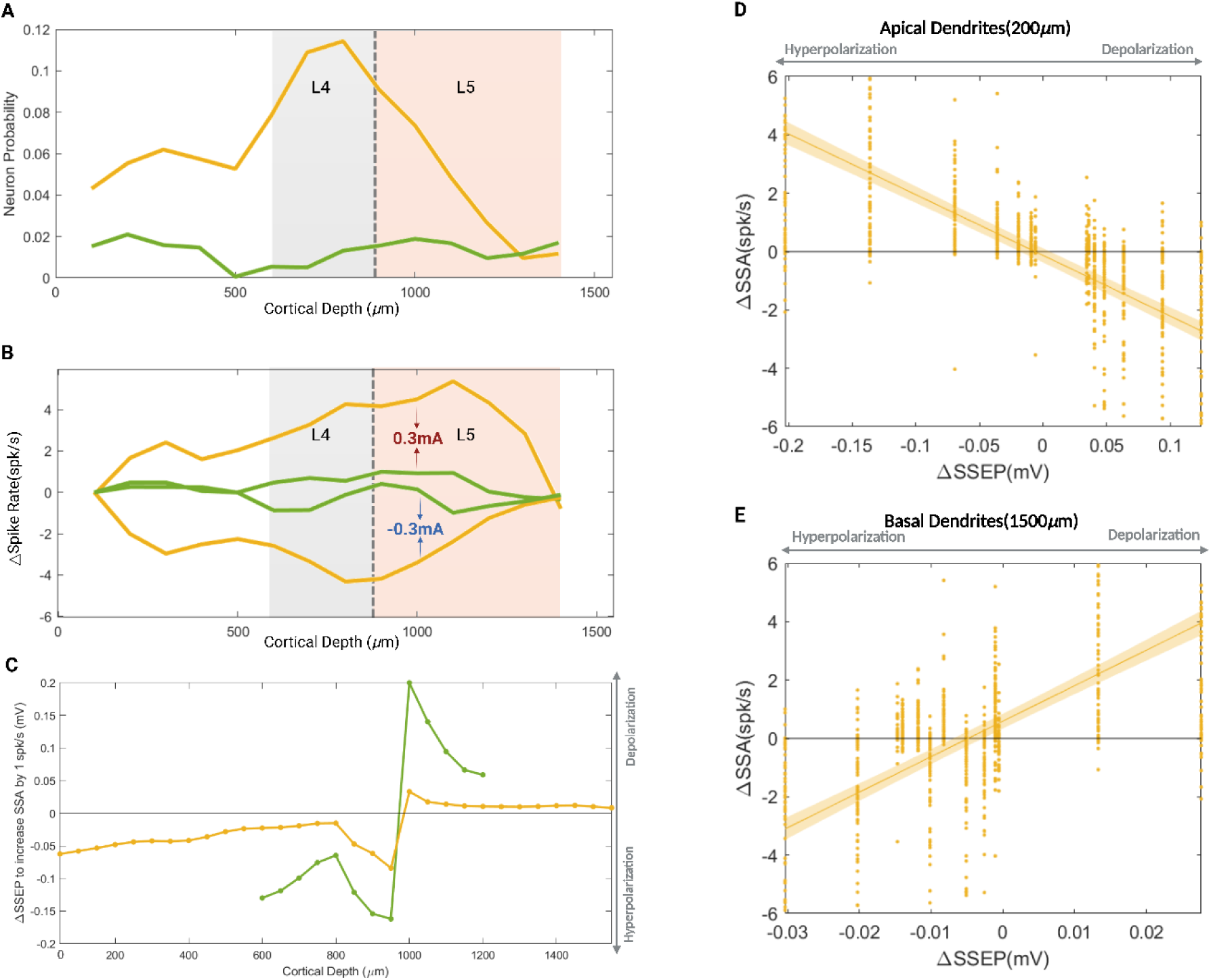
Quantifying the relationship between changes in spontaneous spiking activity and localized membrane polarization. A) Histogram distribution for both RS and FS neuron types binned across different cortical depths. The RS neurons sampled were mostly positioned in layers 4 and 5, while the FS neuron sample was spread approximately equally across all layers. D) The relationship between ΔSSA for RS neurons and ΔSSEP on the apical dendrites was fit with a linear mix effects model (yellow line, shaded area shows 95% confidence intervals). E) Using the same approach, an opposite relationship was found for ΔSSA for RS neurons and ΔSSEP on the basal dendrites. C) The same model was fit using ΔSSEP across all cortical depth. The inverse gradient (i.e. ΔSSEP required to increase SSA by 1 spike/sec) of the fit is shown as a function of cortical depth for RS (yellow) and FS (green) neurons. Note that ΔSSEP effects change polarity at 1000um and that larger changes in ΔSSEP are needed for FS neurons to achieve the same increase in SSA of 1 spike/sec.

Next, we wanted to investigate the relationship between localized changes in membrane polarization as assessed by ΔSSEP (see Fig. 3) and changes SSA. The model shown in Fig. 3 predicts ΔSSEP from different tDCS amplitudes at each cortical depth. Here, we used these model results to define different sets of ΔSSEPs for each RS neuron at each cortical depth (ΔSSEPs_xxxxµm_ i.e. an estimate of the localized membrane polarization experienced by an RS neuron at one specific position on its dendrites or soma, where xxxx represents the specific cortical depth). For each different cortical depth, we then built a different linear mixed effects model using ΔSSEPs_xxxxµm_ to predict ΔSSA (△ SSA ∼ ΔSSEPs_xxxxµm_ +(1|NeuronNumber) +(1|Rat)). Fig 6D and E show the results from the linear models at two cortical depths, 200µm (Fstat=648.8, p<0.001) and 1500µm (Fstat=572.01, p<0.001), selected to represent the opposing effects of tDCS on membrane polarization of apical and basal dendrites of RS (putative pyramidal) neurons. For the apical dendrites, Fig. 6D shows the linear relationship between ΔSSA and ΔSSEP_200µm_. Here, an increase in SSA is associated with a decrease ΔSSEP_200µm_ (i.e. localized hyperpolarization). For the basal dendrites, Fig. 6E shows an opposite, but smaller, effect: i.e. the same increase in ΔSSA is associated with an increase in ΔSSEP_1500µm_ (i.e. localized depolarization). To quantify the effects across all neural compartments, we plotted the inverse gradient of the model fit as a function of cortical depth (see Fig. 6D). Thus, for one RS neuron (yellow line) this Fig. 6D shows the localized changes in ΔSSEP that occur when the neurons SSA is increased by one spike/sec. FS (putative inhibitory) neurons, have much shorter dendrites which do not spread out across multiple cortical layers. Therefore, we next repeated this analysis for FS neurons but across a restricted range of cortical depths (Fig. 6D, green line). Here, we see that for FS neurons, to get the same increase of one spike/sec, much larger changes in ΔSSEP are required.

### 2.6 tDCS effects on evoked spike activity

In addition to tDCS effects on SSA, we analyzed the effects of tDCS on evoked spiking activity. We calculated post-stimulus time histograms (PSTH) for all baseline and tDCS conditions for both RS and FS neurons. The normalized PSTH for the +0.3 mA and -0.3 mA tDCS conditions for RS and FS neurons are shown in Fig 7A-B and E-F respectively. For both RS and FS neurons, we observed that anodic tDCS caused an increase in evoked spiking activity while cathodic tDCS caused a decrease in evoked spiking activity. We then calculated the difference in evoked spiking activity between the baseline and all tDCS conditions for both RS and FS neurons. The data are shown in Fig. 7D and H. These data were fit with a linear mixed-effects model. Fig7. C and G show the linear mixed-effects model fit (solid line) and 95% confidence intervals (shaded) for both RS and FS neuron types respectively. The dots represent the mean Δ evoked spike count data for each tDCS amplitude. We found that tDCS amplitude (Fstat=41.093, p<0.001) had a significant effect on Δ evoked spike count. We also found a significant interaction between tDCS amplitude and neuron type (RS or FS) (Fstat= 22.046, p<0.001) but neuron type alone did not have a significant effect on Δ evoked spike count (Fstat=3.05 p= 0.081). Post hoc testing showed that for RS neurons at anodic 0.1mA (z=4.20, p<0.001), 0.2mA (z=5.14, p<0.001), and 0.3mA (z=4.13, p<0.001) Δ evoked spike counts were significantly higher than baseline. However, at all cathodic amplitudes, there were no significant differences compared to baseline. For FS neurons only at anodic 0.2mA (z=3.01, p=0.0366) was significantly higher than baseline. There were no significant differences compared to the baseline in other amplitude conditions. These results show that tDCS can affect evoked spiking activity. However, in comparison with the effects of tDCS on SSA (see Fig. 5), much stronger electric fields were needed.

**Figure 7.**
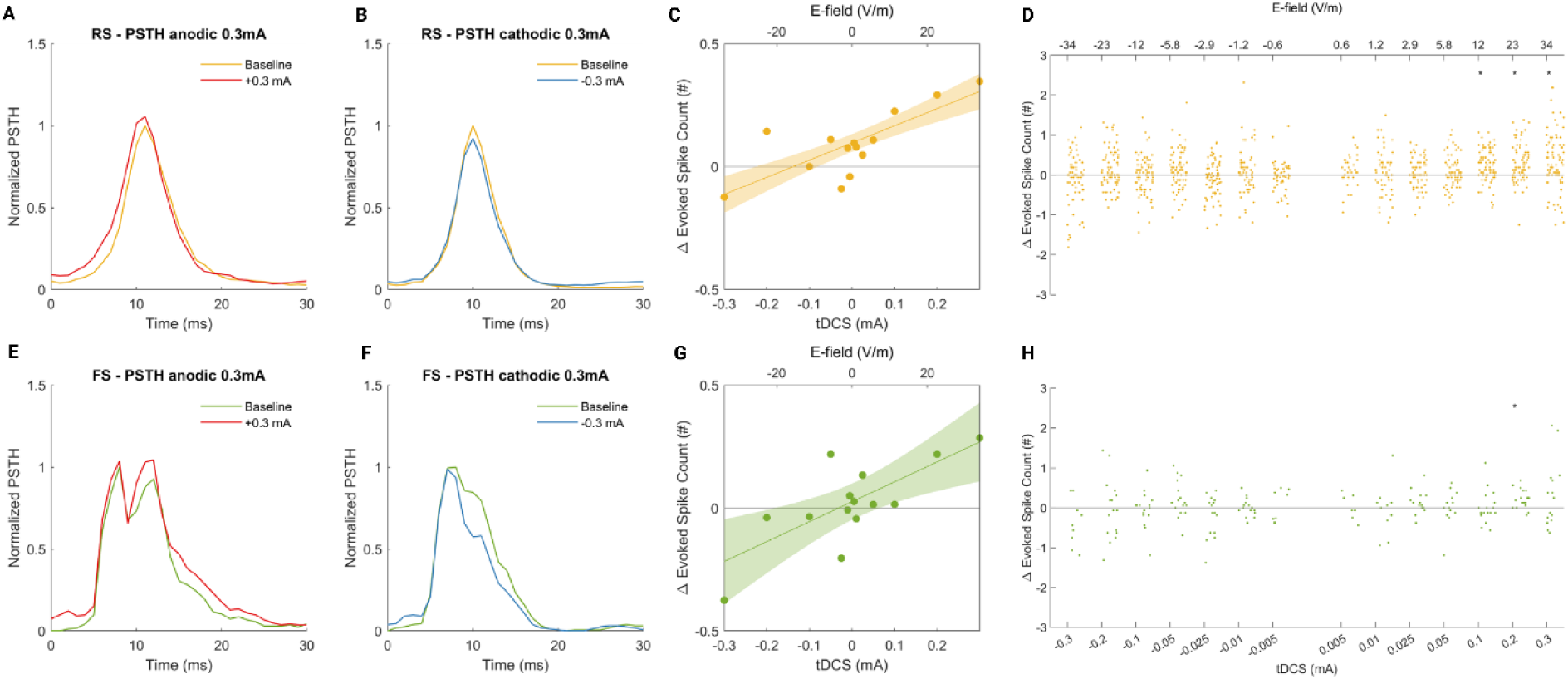
The effect of tDCS on evoked spiking activity using post-stimulus time histograms (PSTH). A) shows normalized PSTH for RS neurons in +0.3mA tDCS condition. B) shows normalized PSTH for RS neurons in -0.3mA tDCS condition. C) shows the linear mixed-effects model fit (solid line) and 95% confidence intervals (shaded) for both RS neurons. The dot represents the mean Δ evoked spike count data for each tDCS amplitude. D) shows the difference in evoked spiking activity between the baseline and all tDCS conditions for both RS neurons. E-H) show the same analysis for FS neurons. The grey dots in D and H represent significant statistical differences.

## 3. Discussion

Previous studies investigating tDCS neural mechanisms have used isolated brain slices with strong electric field strengths that are not achieved in human tDCS ^32–35^. Some of these studies have shown how tDCS can affect membrane polarization ^20,21,36,37^ while others have shown that tDCS works by affecting SSA ^23–25^. However, there is currently a lack of experimental evidence demonstrating how these two proposed tDCS neural mechanisms interact or work in conjunction. Furthermore, it remains unclear how these mechanisms may translate to human tDCS applications, where electric field strengths are always below 1 V/m. In this study, we aimed to partially bridge this gap by developing an in-vivo animal model with realistic electric field strengths. We first used this model to investigate if these proposed tDCS neural mechanisms are also relevant at electric field strengths achieved in human tDCS, and then second to characterize the relationship between tDCS membrane polarization and spontaneous spike rate changes. To do this, we applied tDCS to the rat somatosensory cortex and used a 32-channel silicon probe to measure the electric field strength. Our results demonstrated that a current of 0.005 mA applied to the rat skull generated an electric field of 0.6 V/m at the surface of the somatosensory cortex, approximately equivalent to the electric field achieved in humans when 2 mA tDCS is applied to the scalp ^38^. Applying stronger currents led to stronger electric field strengths with the relationship between skull current amplitude and cortical electric field strength at any particular depth following a linear relationship (Fig. 1).

To investigate the effect of E-field on membrane polarization and cortical excitability we measured SSEPs, in response to foot-shocks, across different cortical layers. Our results demonstrated that the change in SSEP amplitude between the baseline and a tDCS condition (ΔSSEP), showed a complex spatiotemporal pattern across the cortex (Fig. 2). For anodal stimulation we observed an increase in SSEP amplitude in superficial cortical layers, which transitioned to a decrease in SSEP in deeper cortical layers. In other words, we found that SSEP amplitude increased near the apical dendrites and decreased near the soma and basal dendrites during tDCS. In this animal model, the tDCS montage created a radial current flow, parallel to the soma-to-dendritic axis, leading to apical dendrites undergoing hyperpolarization, while somas and basal dendrites undergoing depolarization ^17,18^. Notably, this pattern showed an asymmetrical reversal for cathodal stimulation. This pattern was consistent with the complex pattern of apical dendritic hyperpolarization and soma/basal dendritic depolarization observed for anodal stimulation in pyramidal neurons in brain slices, which also showed an asymmetrical reversal for cathodal stimulation ^39,40^. Thus, here we propose that spatially localized changes in the SSEP serve as a proxy measurement for localized changes in membrane polarization caused by tDCS. Next, we tested if the same pattern of simultaneous hyperpolarization and depolarization was also present at weaker electric field strengths. We found that the changes in skull current and ΔSSEP at any particular cortical depth followed an approximately linear relationship. This showed that the clear pattern of simultaneous hyperpolarization and depolarization observed at stronger electric field strengths scales linearly with current amplitude and is still present at the weak electric field strengths valid for human tDCS (see Fig. 3C and G). A CSD analysis of the SSEP data revealed current sources on the apical dendrites and on the soma/basal dendrites and a current sink in layers 3 and 4, consistent with thalamocortical inputs on pyramidal neurons driving the SSEP ^41,42^ (Fig. 4). This observation also aligns with slice studies which suggest a higher density of thalamocortical boutons in this layer ^43,44^, generating larger excitatory postsynaptic potentials compared to the deeper layer4 and 5, where somas and basal dendrites of most pyramidal neurons are located ^31,45^. Comparing CSD analyses of the baseline and tDCS conditions showed that tDCS had the largest effects on current flow in apical dendrites with changes in current flow also occurring in the soma/basal dendrites ^46^.

Having established that our in-vivo tDCS model caused a complex pattern of hyperpolarization and depolarization in pyramidal neurons, consistent with work in brain slices, we next tested how this would affect SSA. For RS neurons we found a linear relationship between electric field strength and change in SSA (Fig. 5). Post-hoc analysis of our data showed that even weak anodic electric field strengths of 0.6 V/m (at the cortical surface, weaker at deeper depths) caused small but statistically significant changes in spike rate. In contrast, cathodic stimulation required a stronger E-field of -2.9 V/m to cause a significant effect on SSA. This indicates that the anodic tDCS is more effective at modulating SSA in putative pyramidal neurons compared to cathodic tDCS. For the same electric-field strength, FS neurons showed much smaller changes in spike rate. However, these small changes were also linearly related to electric field strength and reversed upon polarity reversal. Brain slice work ^10^ and modeling ^47,48^ have shown that when excitatory and inhibitory neurons are exposed to the same electric field, the elongated morphology of excitatory neurons means they experience a much stronger change in membrane polarization compared to the more spherical inhibitory neurons. Our results in the rat somatosensory cortex are also consistent with a recent publication that has shown similar results in the mouse hippocampus^25^.

One prevailing theory of how tDCS works suggests that it first causes membrane polarization of cortical neurons which in turn leads to changes in SSA ^25^. Computational models of tDCS typically use this assumption to simulate how a DC field can alter the membrane potential and cause changes in SSA. ^21^. Here we present, to the best of our knowledge, the first in-vivo experimental evidence that links changes in membrane polarization to changes in SSA. We showed that anodic tDCS causes a decrease in ΔSSEP (a proxy measurement of membrane depolarization) at the soma which was associated with an increase in SSA for RS neurons and conversely, cathodic tDCS causes an increase in ΔSSEP at the soma, associated with a decrease in SSA for RS neurons. This effect followed a linear relationship with much stronger changes in SSA occurring in RS neurons compared to FS neurons (Fig. 6C). Importantly, we showed that a similar linear relationship exists between ΔSSEP at the basal dendrites and ΔSSA, but that this relationship reverses for ΔSSEP at the apical dendrites. Thus, we present evidence that links the complex pattern of simultaneous hyperpolarization and depolarization caused by tDCS to changes in SSA. This experimental evidence supports the so-called ‘somatic doctrine’ ^14–16^ and shows that this tDCS mechanism is relevant to electric field strengths that occur in human tDCS.

Several studies in human tDCS have discussed the possibility of an inverted u-shaped effect – i.e. the idea that as tDCS amplitude is increased, its effects first increase and then reach a peak before decreasing with higher tDCS amplitudes^49,50^. In our investigation of the electrophysiological mechanisms of tDCS, we found no evidence of a U-shaped effect. All the tDCS mechanisms that we investigated were approximately linearly related to tDCS amplitude and cortical electric field strength, with stronger electric field strengths showing stronger effects. This finding has important implications for human tDCS studies. There is currently a reproducibility crisis in the human tDCS field with several robust studies failing to replicate ^51–55^some of the earlier tDCS studies ^56–59^. On the other hand, as we and others have shown ^25,38^, weak electric fields of less than 1 V/m do have small but measurable effects on membrane polarization, cortical excitability, and SSA. One explanation for the discrepancy between these two fields may be that the small changes in neurophysiological activity observed in animal/brain slice studies at weak electric field strengths do not translate to robust behavioral changes that are measurable in human tDCS studies. Importantly, we showed that all tDCS mechanisms appear to increase with increasing electric field strength. This suggests that novel tDCS approaches that can increase the electric field strength in the brain, such as high amplitude tDCS ^60–62^, multipolar transcranial stimulation ^63^or epicranial stimulation^64^, may lead to more consistent and repeatable effects on human behavior.

Despite the novel insights offered by our in-vivo model of tDCS at realistic electric field strengths, several limitations should be acknowledged. First, our classification of neuronal cell types as regular spiking (RS) and fast-spiking (FS) is an oversimplification that does not fully capture the diversity of interneurons in the cortex. Interneurons play a crucial role in modulating cortical activity, and distinct subtypes may exhibit differential responses to tDCS, as suggested by prior studies ^24,46,47^. Future investigations using advanced classification techniques, such as optogenetic tagging or transgenic mouse lines, which enable the identification of excitatory cells and specific inhibitory cell subtypes, could help clarify the cell type-specific effects of weak electric fields. Secondly, while in our animal model we did demonstrate a linear relationship between electric field strength and neurophysiological effects, the translational relevance of these findings to humans may be more complex. Variability in tDCS outcomes in humans may stem from individual differences in cortical anatomy, electrode positioning, and brain state, factors not accounted for in our animal model. Addressing these limitations in future studies will be crucial for advancing the mechanistic understanding and clinical application of tDCS.

## 4. Method

### 4.1 Animals

A total of 9 male Sprague Dawley rats weighing between 250-400 g (Charles River Laboratories) were used for this study. All animal procedures were approved by the KU Leuven ethics committee for laboratory experimentation (project P072/2020).

### 4.2 Surgery and preparation

Rats were anesthetized with an intraperitoneal injection of urethane (10 mg/mL, Sigma-Aldrich, USA) and placed in a stereotaxic frame (Narishige type SR-6, No. 7905) on a heating pad (ThermoStar Temperature Controller, 69023 pad incl. RWD, China), and their core temperature monitored via a metal rectal probe. We exposed the skull and drilled (US#4 HP 014 drill bit (Meissinger, Germany)) a burr hole relative to bregma at 0 mm AP, -4.00mm ML to target the left primary somatosensory cortex, forelimb region (S1FL).

### 4.3 Electrophysiological recording

For electrophysiology recordings, we used a one-column, 32-channel silicone probe spanning 1550 μm (E32+R-50-S1M-L20 NT, Atlas Neuro, Leuven, Belgium) with a pointy tip allowing insertion without opening the dura. Signals from the probe were amplified (×192), bandpass filtered (0.1 Hz to 7.9 kHz), and digitized (16-bit, 30 kHz) using an RHD 32 Intan head stage (Intan Technologies, Los Angeles, CA) and an Open-Ephys acquisition board (www.open-ephys.org/). The digitized signals were visualized and stored on a PC hard drive using the Open-Ephys GUI v0.6.4. The probe was inserted into the S1FL through the craniotomy and spanned 1550 μm (deepest) to 0 μm (uppermost electrode, where closed to pia matter).

### 4.4 Electrical stimulation setup

To deliver forepaw stimulation a needle electrode (13×0.40mm, Technomed, USA) was inserted into the middle of the right forepaw, and another needle electrode was hooked into the muscle, 1cm from the forepaw. To deliver tDCS we placed a metal disk electrode (1.35mm diameter) with a small amount of conductive gel (Signa Gel, Parker Labs, New Jersey) directly on the skull at 1.5 mm AP, -4.0mm ML. A rectangular metal electrode (1cm2) coated with gel was attached to the tail of the rat as a return. For forelimb stimulation, the forepaw electrode was connected to the positive terminal of a current source STIM1 (AM 2200 analog stimulus isolator, A-M Systems, Sequim, WA), and the electrode in the muscle was connected to the negative terminal. For tDCS the skull electrode was connected to the positive terminal of another current source STIM2 (AM 2200 analog stimulus isolator, A-M Systems, Sequim, WA), and the tail electrode was connected to the negative terminal. Both current sources were controlled via an analog voltage waveform generated on a data acquisition card (100 kHz, NI USB- 6216, National Instruments, Austin, TX).

### 4.4 Experimental protocol

SSEPs were elicited by delivering a single 1mA biphasic pulse (200 µs per phase) every 4 seconds to the forelimb. Data was recorded in 6-minute blocks consisting of a 3-minute baseline condition and a 3- minute tDCS condition. In the tDCS condition, we applied DC stimulation via the skull electrode (3 minutes, 0.005 to 0.3 mA). The SSEP in the baseline and tDCS conditions were then calculated by averaging together the 45 repetitions from the relevant condition. SSEP amplitude on each electrode was quantified as the minimum of the negative trough at around 10ms (either SSEP_baseline_n10_ or SSEP_tDCS_n10_). The change in SSEP amplitude due to tDCS was then calculated as △SSEP_n10_ =SSEP_tDCS_n10_ – SSEP_baseline_n10_.

### 4.5 Extracting single units and spike waveform classification

Note that the Intan amplifier was AC coupled. Thus, in the electrophysiological recording, the DC electrical artifact from skull stimulation was removed by the amplifier hardware filter. The electrical artifact caused by fore-paw stimulation was a single, time-limited pulse, which was removed in software by blanking and linear interpolation. We then performed spike sorting using SpykingCircus and performed manual curation using Phy viewer (https://github.com/cortex-lab/phy). We then extracted spike-times from well-isolated single units and calculated spike rates. To classify spikes as RS or FS we calculated action potential duration as the time between the spike (negative trough) to the first positive peak (see Fig. 5C inset). Spikes with an action potential duration of greater than 0.45 ms were classified as RS, while spikes with a duration of less than or equal to 0.45 ms were classified as FS.

### 4.6 Measurement and calculation of electric field strength

The amplifier in the RHD 32 Intan headstage used in the electrophysiology setup is AC coupled and thus cannot measure the electric field caused by direct current. Therefore, to measure the electric field caused by our transcranial stimulation setup we delivered a 100Hz sine wave through the skull electrode, 1-minute duration, at 0.01, 0.015, 0.02, and 0.025 mA. These amplitudes were chosen to be low enough to ensure that they did not saturate the amplifier which had an input range of +/- 5mV.

After the experiment, we then performed a discrete Fourier transform of the recorded data and extracted the voltage amplitude of the 100Hz component across each of the 32 channels for each of the delivered stimulation amplitudes. An example from one animal example shown in Fig. 1A. This was procedure was repeated in 8 animals (data not collected in one animal). We then pooled the data from 8 animals and a fit it with a linear mixed-effects model with the slope of the linear model being fit across the different voltages (i.e. stimulation amplitude was used to predict the measured voltage, fit to all data shown in Fig. 1B). The model notation was Voltage∼tDCS*Depth+(1|Rat). We then calculated the electric field by taking the spatial derivative of the modeled voltage data (Fig. 1C). The linear fit could then be extrapolated to predict the electric field strengths that would present a higher tDCS amplitude (Fig. 1D), which could not be measured directly because of amplifier saturation.

### 4.7 Current source analysis

Calculation of the CSD for the baseline and tDCS conditions, for all different stimulation amplitudes, followed the same procedure: First, the SSEPs for that condition for each of the 9 rats were averaged together. For one time point in the SSEP, we then calculated the second spatial derivative along the depth axis and applied a 3-point running average (i.e. a span of 3 electrodes) to smooth. This was repeated for each time point in the SSEP yielding the full CSD (Fig. 4)^65,66^.

### 4.8 Statistics

The data from all 9 rats were pooled and a linear mixed-effects model was used to quantify the effect of tDCS amplitude (tDCS) and cortical depth (Depth) on △SSEP_n10_, with tDCS and depth being fixed effects and rat a random effect: △SSEP∼tDCS*Depth+(1|Rat). ANOVA was then performed on the model output to test the significance of the fixed effect.

Spiking data from all nine rats were pooled and the difference in spontaneous spike activity between the baseline and tDCS condition was calculated (△SSA). As described above the neuron type was defined as either RS or FS based on action potential duration. We then used a linear mixed effects model to quantify the effect of tDCS amplitude (tDCS) and neuron type on △SSA, with tDCS and neuron type being fixed effects and rat and neuron number being random effects: △ SSA ∼tDCS*NeuronType+(1|NeuronNumber) +(1|Rat). This approach follows best statistical practice for spiking data^67^. ANOVA was then performed on the model output to test the significance of the fixed effect. Post-hoc testing was conducted using the two-sided Wilcoxon signed rank to test if the △SSA in each condition was significantly different than zero. All reported p-values were Bonferroni corrected for 14 multiple comparisons (i.e. all the tDCS amplitude conditions). The alpha of all tests was set to 0.05. The same approach was used for the effect of tDCS amplitude and neuron type on evoked spiking activity.

## Acknowledgments

This work was supported by FWO Funding G0B4520N and NIH Funding 1R01MH123508-01. Liyi Chen was funded by the China Scholarship Council at CSC202006380043.

## Author contributions

Liyi Chen designed the experiments, performed all experiments, processed and analyzed data, and wrote the manuscript. Alireza Majdi assisted in experiments and data collection. Boateng Asamoah supervised the progress of the project and refined the manuscript. Myles McLaughlin: designed the experiment, supervised and coordinated the research, visualized data, and approved the final manuscript.

## Reference

1. Chase, H. W., Boudewyn, M. A., Cameron, •, Carter, S. & Phillips, M. L. Transcranial direct current stimulation: a roadmap for research, from mechanism of action to clinical implementation. Mol Psychiatry 25, 397–407 (2020).

2. Flöel, A. tDCS-enhanced motor and cognitive function in neurological diseases. Neuroimage 85, 934–947 (2014).

3. Pisoni, A. et al. Cognitive Enhancement Induced by Anodal tDCS Drives Circuit-Specific Cortical Plasticity. Cerebral Cortex 28, 1132–1140 (2018).

4. Chan, M. M. Y., Yau, S. S. Y. & Han, Y. M. Y. The neurobiology of prefrontal transcranial direct current stimulation (tDCS) in promoting brain plasticity: A systematic review and meta-analyses of human and rodent studies. Neurosci Biobehav Rev 125, 392–416 (2021).

5. Jonker, Z. D. et al. No effect of anodal tDCS on motor cortical excitability and no evidence for responders in a large double-blind placebo-controlled trial. Brain Stimul 14, 100–109 (2021).

6. Horvath, J. C., Forte, J. D. & Carter, O. Quantitative Review Finds No Evidence of Cognitive Effects in Healthy Populations From Single-session Transcranial Direct Current Stimulation (tDCS). Brain Stimul 8, 535–550 (2015).

7. Galli, G., Vadillo, M. A., Sirota, M., Feurra, M. & Medvedeva, A. A systematic review and meta-analysis of the effects of transcranial direct current stimulation (tDCS) on episodic memory. Brain Stimul 12, 231–241 (2019).

8. Nikolin, S., Martin, D., Loo, C. K. & Boonstra, T. W. Effects of TDCS dosage on working memory in healthy participants. Brain Stimul 11, 518–527 (2018).

9. Chase, H. W., Boudewyn, M. A., Carter, C. S. & Phillips, M. L. Transcranial direct current stimulation: a roadmap for research, from mechanism of action to clinical implementation. Mol Psychiatry 25, 397–407 (2020).

10. Radman, T., Ramos, R. L., Brumberg, J. C. & Bikson, M. Role of cortical cell type and morphology in subthreshold and suprathreshold uniform electric field stimulation in vitro. Brain Stimul 2, (2009).

11. Lafon, B., Rahman, A., Bikson, M. & Parra, L. C. Direct Current Stimulation Alters Neuronal Input/Output Function. Brain Stimul 10, 36–45 (2017).

12. Bikson, M. et al. Effect of uniform extracellular DC electric fields on excitability in rat hippocampal slices in vitro. Journal of Physiology 557, 175–190 (2004).

13. Kronberg, G., Bridi, M., Abel, T., Bikson, M. & Parra, L. C. Direct Current Stimulation Modulates LTP and LTD: Activity Dependence and Dendritic Effects. Brain Stimul 10, 51–58 (2017).

14. Nitsche, M. A. & Paulus, W. Excitability changes induced in the human motor cortex by weak transcranial direct current stimulation. J Physiol 527, 633 (2000).

15. Gorman, A. L. Differential patterns of activation of the pyramidal system elicited by surface anodal and cathodal cortical stimulation. J Neurophysiol 29, 547–564 (1966).

16. D, Purpura. P. & Mcmurtry, J. G. INTRACELLULAR ACTIVITIES AND EVOKED POTENTIAL CHANGES DURING POLARIZATION OF MOTOR CORTEX. J Neurophysiol 28, 166–185 (1965).

17. Rahman, A. et al. Cellular effects of acute direct current stimulation: somatic and synaptic terminal effects. J Physiol 591, 2563–2578 (2013).

18. Radman, T., Ramos, R. L., Brumberg, J. C. & Bikson, M. Role of cortical cell type and morphology in subthreshold and suprathreshold uniform electric field stimulation in vitro. Brain Stimul 2, (2009).

19. Lafon, B., Rahman, A., Bikson, M. & Parra, L. C. Direct Current Stimulation Alters Neuronal Input/Output Function. Brain Stimul 10, 36–45 (2017).

20. Jackson, M. P. et al. Animal models of transcranial direct current stimulation: Methods and mechanisms. Clinical Neurophysiology 127, 3425–3454 (2016).

21. Molaee-Ardekani, B. et al. Effects of transcranial Direct Current Stimulation (tDCS) on cortical activity: A computational modeling study. Brain Stimul 6, 25–39 (2013).

22. Stagg, C. J., Antal, A. & Nitsche, M. A. Physiology of Transcranial Direct Current Stimulation. Journal of ECT 34, 144–152 (2018).

23. Ozen, S. et al. Transcranial Electric Stimulation Entrains Cortical Neuronal Populations in Rats. Journal of Neuroscience 30, 11476–11485 (2010).

24. Vöröslakos, M. et al. Direct effects of transcranial electric stimulation on brain circuits in rats and humans. Nat Commun 9, (2018).

25. Farahani, F., Khadka, N., Parra, L. C., Bikson, M. & Vöröslakos, M. Transcranial electric stimulation modulates firing rate at clinically relevant intensities. Brain Stimul 17, 561–571 (2024).

26. Milighetti, S. et al. Effects of tDCS on spontaneous spike activity in a healthy ambulatory rat model. Brain Stimul 13, 1566–1576 (2020).

27. Liu, A. et al. Immediate neurophysiological effects of transcranial electrical stimulation. Nature Communications 2018 9:1 9, 1–12 (2018).

28. Khatoun, A., Asamoah, B. & Laughlin, M. M. Investigating the feasibility of epicranial cortical stimulation using concentric-ring electrodes: A novel minimally invasive neuromodulation method. Front Neurosci 13, 440965 (2019).

29. Rahman, A. et al. Cellular effects of acute direct current stimulation: somatic and synaptic terminal effects. J Physiol 591, 2563–2578 (2013).

30. Meyer, H. S. et al. Cell Type–Specific Thalamic Innervation in a Column of Rat Vibrissal Cortex. Cerebral Cortex 20, 2287–2303 (2010).

31. Reyes-Puerta, V., Sun, J. J., Kim, S., Kilb, W. & Luhmann, H. J. Laminar and Columnar Structure of Sensory-Evoked Multineuronal Spike Sequences in Adult Rat Barrel Cortex In Vivo. Cereb Cortex 25, 2001–2021 (2015).

32. Bikson, M. et al. Effects of uniform extracellular DC electric fields on excitability in rat hippocampal slices in vitro. J Physiol 557, 175–190 (2004).

33. Chan, C. Y. & Nicholson, C. Modulation by applied electric fields of Purkinje and stellate cell activity in the isolated turtle cerebellum. J Physiol 371, 89–114 (1986).

34. Rahman, A., Lafon, B., Parra, L. C. & Bikson, M. Direct current stimulation boosts synaptic gain and cooperativity in vitro. J Physiol 595, 3535–3547 (2017).

35. Reato, D., Bikson, M. & Parra, L. C. Lasting modulation of in vitro oscillatory activity with weak direct current stimulation. J Neurophysiol 113, 1334–1341 (2015).

36. Jamil, A. & Nitsche, M. A. What Effect Does tDCS Have on the Brain? Basic Physiology of tDCS. Curr Behav Neurosci Rep 4, 331–340 (2017).

37. Modolo, J., Denoyer, Y., Wendling, F. & Benquet, P. Physiological effects of low-magnitude electric fields on brain activity: advances from in vitro, in vivo and in silico models. Curr Opin Biomed Eng 8, 38 (2018).

38. Huang, Y. et al. Measurements and models of electric fields in the in vivo human brain during transcranial electric stimulation. Elife 6, (2017).

39. Farahani, F., Khadka, N., Parra, L. C., Bikson, M. & Vöröslakos, M. Transcranial electric stimulation modulates firing rate at clinically relevant intensities. Brain Stimul 17, 561–571 (2024).

40. Tremblay, R., Lee, S. & Rudy, B. GABAergic interneurons in the neocortex: From cellular properties to circuits. Neuron 91, 260 (2016).

41. Jacobs, K. M. Somatosensory System. Encyclopedia of Clinical Neuropsychology 1–7 (2018) doi:10.1007/978-3-319-56782-2_359-2.

42. O’Reilly, C., Iavarone, E., Yi, J. & Hill, S. L. Rodent somatosensory thalamocortical circuitry: Neurons, synapses, and connectivity. Neurosci Biobehav Rev 126, 213–235 (2021).

43. Smith, P. H., Uhlrich, D. J., Manning, K. A. & Banks, M. I. Thalamocortical projections to rat auditory cortex from the ventral and dorsal divisions of the medial geniculate nucleus. Journal of Comparative Neurology 520, 34–51 (2012).

44. Ji, X. Y. et al. Thalamocortical Innervation Pattern in Mouse Auditory and Visual Cortex: Laminar and Cell-Type Specificity. Cerebral Cortex 26, 2612–2625 (2016).

45. Meyer, H. S. et al. Cell Type–Specific Thalamic Innervation in a Column of Rat Vibrissal Cortex. Cerebral Cortex 20, 2287–2303 (2010).

46. An, S., Kilb, W. & Luhmann, H. J. Sensory-Evoked and Spontaneous Gamma and Spindle Bursts in Neonatal Rat Motor Cortex. The Journal of Neuroscience 34, 10870 (2014).

47. Ye, H. & Steiger, A. Neuron matters: electric activation of neuronal tissue is dependent on the interaction between the neuron and the electric field. J Neuroeng Rehabil 12, 1–9 (2015).

48. Kepecs, A. & Fishell, G. Interneuron cell types are fit to function. Nature 2014 505:7483 505, 318–326 (2014).

49. Monte-Silva, K. et al. Dose-Dependent Inverted U-Shaped Effect of Dopamine (D2-Like) Receptor Activation on Focal and Nonfocal Plasticity in Humans. The Journal of Neuroscience 29, 6124 (2009).

50. Ehrhardt, S. E., Filmer, H. L., Wards, Y., Mattingley, J. B. & Dux, P. E. The influence of tDCS intensity on decision-making training and transfer outcomes. J Neurophysiol 125, 385–397 (2021).

51. Westwood, S. J., Olson, A., Miall, R. C., Nappo, R. & Romani, C. Limits to tDCS effects in language: Failures to modulate word production in healthy participants with frontal or temporal tDCS. Cortex 86, 64–82 (2017).

52. Horvath, J. C., Forte, J. D. & Carter, O. Quantitative Review Finds No Evidence of Cognitive Effects in Healthy Populations From Single-session Transcranial Direct Current Stimulation (tDCS). Brain Stimul 8, 535–550 (2015).

53. Horvath, J. C., Forte, J. D. & Carter, O. Evidence that transcranial direct current stimulation (tDCS) generates little-to-no reliable neurophysiologic effect beyond MEP amplitude modulation in healthy human subjects: A systematic review. Neuropsychologia 66, 213–236 (2015).

54. Mancuso, L. E., Ilieva, I. P., Hamilton, R. H. & Farah, M. J. Does Transcranial Direct Current Stimulation Improve Healthy Working Memory?: A Meta-analytic Review. J Cogn Neurosci 28, 1063–1089 (2016).

55. Buch, E. R. et al. Effects of tDCS on motor learning and memory formation: A consensus and critical position paper. Clinical Neurophysiology 128, 589–603 (2017).

56. Zimerman, M. et al. Neuroenhancement of the aging brain: Restoring skill acquisition in old subjects. Ann Neurol 73, 10–15 (2013).

57. Nitsche, M. A. et al. Facilitation of implicit motor learning by weak transcranial direct current stimulation of the primary motor cortex in the human. J Cogn Neurosci 15, 619–626 (2003).

58. Dedoncker, J., Brunoni, A. R., Baeken, C. & Vanderhasselt, M. A. A Systematic Review and Meta-Analysis of the Effects of Transcranial Direct Current Stimulation (tDCS) Over the Dorsolateral Prefrontal Cortex in Healthy and Neuropsychiatric Samples: Influence of Stimulation Parameters. Brain Stimul 9, 501–517 (2016).

59. Hummel, F. C. et al. Facilitating skilled right hand motor function in older subjects by anodal polarization over the left primary motor cortex. Neurobiol Aging 31, 2160–2168 (2010).

60. El Jamal, C., et al. Tolerability and blinding of high-definition transcranial direct current stimulation among older adults at intensities of up to 4 mA per electrode. Brain Stimul 16, 1328–1335 (2023).

61. Reckow, J. et al. Tolerability and blinding of 4×1 high-definition transcranial direct current stimulation (HD-tDCS) at two and three milliamps. Brain Stimul 11, 991–997 (2018).

62. Khadka, N. et al. Adaptive current tDCS up to 4 mA. Brain Stimul 13, 69–79 (2020).

63. Grossman, N. et al. Noninvasive Deep Brain Stimulation via Temporally Interfering Electric Fields. Cell 169, 1029–1041.e16 (2017).

64. Khatoun, A., Asamoah, B. & Laughlin, M. M. Investigating the feasibility of epicranial cortical stimulation using concentric-ring electrodes: A novel minimally invasive neuromodulation method. Front Neurosci 13, 440965 (2019).

65. Nicholson, C. & Freeman, J. A. Theory of current source-density analysis and determination of conductivity tensor for anuran cerebellum. J Neurophysiol 38, 356–368 (1975).

66. Freeman, J. A. & Nicholson, C. Experimental optimization of current source-density technique for anuran cerebellum. J Neurophysiol 38, 369–382 (1975).

67. Yu, Z. et al. Beyond t test and ANOVA: applications of mixed-effects models for more rigorous statistical analysis in neuroscience research. Neuron 110, 21–35 (2022).

